# Physiologically distinct neurons within the ventrolateral periaqueductal gray are not defined by mu-opioid receptor expression but are differentially activated by persistent inflammation

**DOI:** 10.1101/2021.06.16.448597

**Authors:** Kylie B. McPherson, Courtney A. Bouchet, Susan L. Ingram

## Abstract

The ventrolateral periaqueductal gray (vlPAG) is a key structure within the descending pain modulatory pathway and an important target for opioid-induced analgesia. This area contains heterogeneous neurons with respect to neurotransmitter and receptor expression so it is difficult to define vlPAG neurons that contribute to pain and analgesia. Characterization of intrinsic membrane properties of 371 vlPAG neurons from female and male Long-Evans rats identified 4 neuron types with distinct intrinsic firing patterns: *Phasic, Tonic, Onset*, and *Random. Phasic* and *Tonic* neurons comprise the majority of the neurons sampled. Mu-opioid receptor (MOR) expression was determined by the ability of the selective MOR agonist DAMGO to activate G protein-coupled inwardly-rectifying potassium channel (GIRK) currents. Opioid-sensitive and -insensitive neurons were observed within each neuron type in naïve rats and in rats pretreated with Complete Freund’s adjuvant in a hindpaw to produce persistent inflammation. The presence of low threshold spikes (LTS) did not correlate with MOR-mediated GIRK currents indicating that MOR expression alone does not define a physiologically distinct neuron type in the vlPAG. MOR activation inhibited firing in nearly all spontaneously active neurons, both in naïve and persistent inflammation conditions. CFA-induced inflammation increased Fos expression at both acute (2 h) and persistent inflammation (5-7 d) time points. However, persistent, but not acute, inflammation selectively enhanced spontaneous firing and lowered firing thresholds of *Phasic* neurons which was maintained in the absence of synaptic inputs. Taken together, persistent inflammation selectively activates *Phasic* neurons, of which only a subset was opioid-sensitive.

**Significance statement:** Intrinsic membrane properties define separate vlPAG neurons types that are functionally important. Persistent, and not acute, inflammation selectively activates *Phasic* firing vlPAG neurons that are not defined by MOR expression. Although the vlPAG is known to contribute to the descending *inhibition* of pain, the activation of a single physiologically-defined neuron type in the presence of inflammation may represent a mechanism by which the vlPAG participates in descending *facilitation* of pain.

## Introduction

The ventrolateral periaqueductal gray (vlPAG) is an important site contributing to the descending modulation of pain (Bodnar, 2000; Heinricher and Ingram, 2020). Electrical and chemical stimulation of the vlPAG induces strong antinociception in animals (Reynolds, 1969; Mayer and Liebeskind, 1974; Soper and Melzack, 1982) and pain relief in humans (Hosobuchi et al., 1977; Barbaro, 1988; Bittar et al., 2005). Direct opioid microinjections into the vlPAG induce antinociception defining the vlPAG as an area involved in pain inhibition (Yaksh et al., 1976; Jensen and Yaksh, 1989; Morgan et al., 2020). The vlPAG neurons express many different neurotransmitters, neuropeptides and receptors (Bagley and Ingram, 2020). Current genetic strategies attempt to define circuits by targeting expression of specific markers, such as the mu-opioid receptor (MOR), but the heterogeneity within the vlPAG makes genetic strategies difficult to interpret. In our studies, intrinsic membrane properties of vlPAG neurons were examined to define functional populations of neurons that are modulated by opioids and pain.

Opioids disinhibit vlPAG output neurons to the rostral ventromedial medulla (RVM) to produce analgesia (Basbaum and Fields, 1984; Cheng et al., 1986; Lau et al., 2020). In a subpopulation of opioid-sensitive vlPAG neurons, postsynaptic MORs activate G protein-coupled inwardly-rectifying potassium channels (GIRKs) hyperpolarizing the neurons (Barbaresi and Manfrini, 1988; Chieng and Christie, 1994; Kalyuzhny and Wessendorf, 1998; McPherson et al., 2018), some of which are local GABAergic neurons (Basbaum and Fields, 1978; Barbaresi and Manfrini, 1988; Reichling and Basbaum, 1990; Kalyuzhny and Wessendorf, 1998). Although it is tempting to use MOR as a cell-type specific marker, morphological features of opioid-sensitive and -insensitive neurons are similar (Chieng and Christie, 1994) and MORs are expressed on both GABAergic and non-GABA neurons within this region (Zhang et al., 2020). Further, electrophsyiological characteristics that have been used to identify MOR-expressing neurons, have been limited to small numbers of vlPAG neurons (Park et al., 2010; Lau et al., 2020). Thus, a more comprehensive study was necessary to determine if opioid-sensitive neurons represent a single functional subset of neurons in the vlPAG with distinct firing properties.

Noxious stimuli excite a subpopulation of vlPAG neurons (Heinricher et al., 1987; Tryon et al., 2016) and promote Fos expression within the vlPAG (Keay and Bandler, 1993; Keay et al., 2000). Chronic pain activates *in vivo* vlPAG activity (Samineni et al., 2017) and increased BOLD responses in the PAG are observed in humans experiencing secondary hyperalgesia (Zambreanu et al., 2005), indicating activation of PAG neurons in pain states. The intrinsic membrane and firing properties of vlPAG neurons that are activated by noxious stimuli have not been characterized. Thus, the purpose of this study was to characterize the intrinsic membrane properties of vlPAG neurons to determine whether specific subpopulations are sensitive to opioids or selectively activated by acute or persistent CFA-induced inflammation.

## Methods

### Animals

Female and male Long Evans rats (Harlan Laboratories and bred in house; 30-90 d postnatal for electrophysiology) were group housed with environmental enrichment squares and toy bones. Lights were on a 12 h light and dark cycle, with food and water provided *ad libitum*. All experiments were approved by the Animal Care and Use Committee at Oregon Health & Science University. A subset of the studies was done in Sprague Dawley rats (n = 70) and observed no strain differences.

### CFA treatment

Rats were lightly anesthetized with isoflurane and 100 µL of Complete Freund’s adjuvant (CFA; Sigma) was injected into the left hindpaw. Rats were used 2 h or 5-7 d following injections (6 d for Fos immunohistochemistry).

### Fos immunohistochemistry

Rats were deeply anesthetized with isoflurane and perfused with phosphate-buffered saline (PBS), followed by 4% paraformaldehyde (PFA) either 2 h or 6 d after CFA injection in the hind paw. The brains were post-fixed for an additional 24 hrs before being transferred and incubated in 30% sucrose in PBS at 4 °C for 2-3 days. Brains were then frozen in dry ice and stored at 80 °C until coronal brain slices (40 µM) were cut using a Leica CM2050 S cryostat. Sections were washed in PBS, blocked in 3% normal goat serum in PBS with 0.25% Triton X-100 (PBS-Tx), and incubated for 24 h at 4 °C with anti-Fos antibody (1:8000 dilution; Cell Signaling Technology, catalog #2250) in blocking solution. Sections were then washed in PBS and incubated with biotinylated goat anti-rabbit secondary antibody (1:600 dilution; Vector Laboratories, catalog #BA-1000) in PBS-Tx and 1% NGS for 2 h at room temperature. After another wash in PBS, sections were incubated in avidin-biotin-peroxidase complex (ABC Elite kit, PK-6100; Vector Laboratories) in PBS containing 0.5% Triton X-100. Lastly, sections were developed in 3,3-diaminobenzidine (DAB) for 3-5 m (until uniform vlPAG staining occurred with minimal increase in background staining). Sections were then mounted onto chromalumgelatin-coated slides. After drying the slides were dehydrated using a progression of alcohol (30%, 60%, 90%, 100%, 100% ethanol) followed by Citrasolv (Fisher Scientific) before coverslipping with Permount (Sigma Aldrich). Bright-field images of immunoreactive neurons in the vlPAG were captured digitally using an Olympus BX51 fluorescence microscope at 10x magnification. Regions of interest within the vlPAG were measured and Fos labeled neurons were manually quantified by an investigator blind to the treatment conditions, to give a Fos+ nuclei/mm^2^ value. The 4 sections per rat were averaged so that each rat represented an n = 1 for staining across the rostral-caudal extent of the ventrolateral column.

### Electrophysiology

Rats were anesthetized with isoflurane and brains were quickly removed and immersed in ice-cold sucrose aCSF containing the following (in mM): 75 NaCl, 2.5 KCl, 0.1 CaCl_2_, 6 MgSO_4_, 1.2 NaH_2_PO_4_, 25 NaHCO_3_, 2.5 dextrose, 50 sucrose. Coronal slices containing the vlPAG were cut 220 µm thick with a vibratome (Leica Microsystems). Slices were placed in a holding chamber oxygenated with aCSF containing the following (in mM): 126 NaCl, 21.4 NaHCO_3_, 11.1 dextrose, 2.5 KCl, 2.4 CaCl_2_, 1.2 MgCl_2_, and 1.2 NaH_2_PO_4_, and equilibrated with 95% O_2_ /5% CO_2_ at 32 °C until recording. Brain slices were placed into a recording chamber on an upright Zeiss Examiner Z1 and superfused with 32 °C aCSF. Whole-cell recordings were conducted with pipettes (3-4 MΩ) with KMeSO_4_ internal solution containing the following (in mM): 138 CH_3_KO_4_S, 10 HEPES, 10 KCl, 1 MgCl_2_, 1 EGTA, 0.3 CaCl_2_, 4 MgATP, and 3 NaGTP, pH 7.3-7.4. A junction potential of 15 mV was corrected at the beginning of the experiments. Access was monitored throughout the experiments. During whole-cell voltage-clamp recordings, neurons were held at -70 mV. During current-clamp recordings, no holding current was used. Data acquisition was completed with Axopatch 200B Microelectrode Amplifier (Molecular Devices) at 5 kHz and low-pass filtered at 2 kHz. Currents were digitized with InstruTECH ITC-18 (HEKA), collected via AxoGraph Data Acquisition software and analyzed using AxoGraph (AxoGraph Scientific). All analyses were blind to treatment.

### Neuron type characterization

We recorded from 223 neurons in the vlPAG from naïve and 148 neurons from CFA-treated male and female Long-Evans rats. A subset of neurons (n = 59) from naïve rats recorded over the same time period as the recordings from acute and chronic CFA-treated rats were used for naïve vs CFA comparisons to control for handling, housing stress, etc. Only neurons with stable resting membrane potentials (RMP) exhibiting action potentials that crossed 0 mV when depolarized by current step protocols were used for electrophysiological analysis. In current-clamp, 2 s long depolarizing steps from 0 pA to 120 pA in 20 pA increments, with a 1s delay between steps were used to evaluate firing patterns of vlPAG neurons. Spontaneous firing frequency was determined using the total number of action potentials in a 3 s trace. Firing frequency during the current steps was determined by the total number of action potentials divided by the time spent firing. Low threshold spiking (LTS) was determined using a 500 ms long hyperpolarizing current step (−50 pA), which either did or did not elicit a spike upon the offset of the current step.

*Input resistance (R*_*in*_) was determined by calculating the steady state current response from a -10 mV hyperpolarizing step in voltage-clamp. *Resting membrane potential (RMP)* was determined by averaging the baseline traces prior to the series of depolarizing current steps. RMP was corrected for the junction potential for the KMeSO_4_ intracellular solution (−15 mV). *Capacitance* values were calculated from a -10 mV hyperpolarizing voltage step using Axograph X.

### GIRK currents

GIRK current responses to DAMGO were measured in voltage-clamp after completing the current-clamp protocols used to assess firing patterns. Neurons were held at -70 mV in voltage-clamp. After 2-3 mins of a stable baseline, a maximal concentration of selective MOR agonist DAMGO (5 µM) was superfused over the slice until the outward current reached the peak. The nonselective opioid receptor antagonist naloxone (10 µM) was then superfused until a steady return baseline was achieved. GIRK currents were measured as the difference between the peak of the current compared to an average of currents measured at baseline and reversal in the presence of naloxone. A neuron was considered opioid-insensitive if there was no change in current induced by the agonist. GIRK current traces were analyzed blind to the neuron type and treatment.

### Loose patch recording

Pipettes with KMeSO_4_ internal solution (2-3 MΩ) were used to get a cell-attached loose patch seal (Beckstead et al., 2004) to passively measure firing rates of intact, spontaneously active vlPAG neurons without altering intracellular mileau. After 2-3 mins of a stable baseline, the MOR agonist DAMGO (5 µM) was superfused over the slice for 4-5 min followed by antagonist naloxone (10 µM). Traces were excluded from the data set if they had a run up or run down of firing activity throughout the entire recording.

### Experimental Design and Statistical Analysis

All data are expressed as mean ± SEM. Data were analyzed with Prism 9 (GraphPad Software). Each electrophysiological recording from a single neuron is treated as an individual observation; however, all data sets contain recordings from at least 8 different animals. Differences between groups were assessed using ANOVA and t-tests when appropriate (significance denoted as *p < 0.05, **p < 0.01, ***p < 0.001, and ****p < 0.0001).

## Results

### vlPAG neurons can be divided into 4 neuron types with distinct firing properties

The firing patterns and associated membrane properties of vlPAG neurons from Long-Evans rats were evaluated and split into 4 neuron types based on distinct firing patterns: *Phasic, Tonic, Onset*, and *Random*. No notable sex differences in firing pattern or firing threshold were observed, subsequently the recordings from female and male rats were combined for analysis. Of the 223 neurons in naïve rats, 103 displayed firing patterns where firing duration within each step decreased as the stimulus intensity increased (*Phasic*, Fig. 1). The second largest group (77/223) fired throughout each of the depolarizing steps above 20 pA (*Tonic*, Fig. 1). Approximately 10% of the neurons fired 1-2 action potentials at the beginning of each depolarizing step (*Onset*; Fig. 1) and another ∼10% displayed randomly timed action potentials throughout (*Random*, Fig. 1). All neurons were evaluated for intrinsic membrane properties including resting membrane potential (RMP), membrane resistance (R_in_), capacitance, and spontaneous firing activity (Table 1). The topographical distribution was evaluated for a large subset of neurons showing no unique organization or clustering of the different types throughout the rostral-caudal axis of the vlPAG (n = 117, chi-square, p = 0.2). Previous work in the lab has used Sprague Dawley rats, so we also examined firing patterns from this strain and observed the same neuron types in similar proportions: 57% *Phasic* (25/44), 18% *Tonic* (8/44), 14% *Random* (6/44), and 11% *Onset* (5/44).

**Figure 1.**
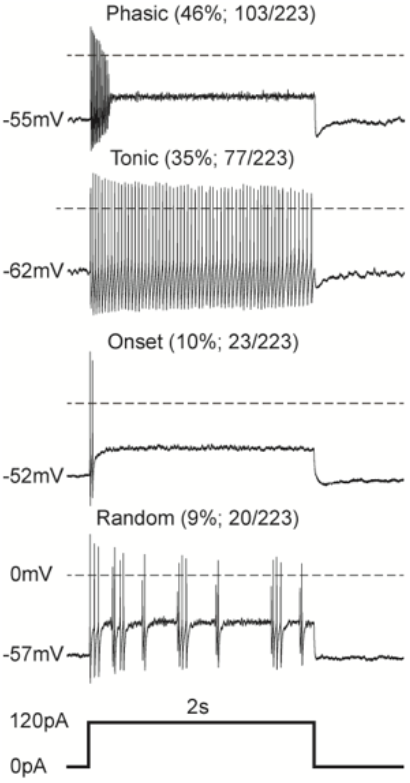
The vlPAG contains 4 neuron types defined by distinct firing patterns. Representative traces from vlPAG neurons from naïve rats showing 4 distinct neuron types based on their response to a 2 s long 120 pA depolarizing current step. The proportion of neurons exhibiting the firing pattern out of the total neurons recorded in naïve rats and the resulting percentage are represented in parentheses. Resting membrane potential is noted at the beginning of each trace and the dotted line represents 0 mV.

**Table 1.** Electrophysiological properties of 4 neuron types in naïve rats. The number of neurons recorded: *Phasic* (n=103), *Tonic* (n=77), *Onset* (n=23), and *Random* (n=20). Resting membrane potential (F_(3, 219)_ = 19.1, ****p < 0.0001; Tukey’s multiple comparisons; *Phasic* vs. *Tonic* and *Tonic* vs. *Onset*, p < 0.0001, *Onset* vs. *Random*, p = 0.04), Membrane resistance (F_(3, 219)_ = 3.6, *p = 0.01; Tukey’s multiple comparisons test; *Phasic* vs. *Random*, p = 0.02), and Capacitance (F_(3, 219)_ = 18.4, ****p < 0.0001; Tukey’s multiple comparisons test; *Phasic* vs. *Tonic*, ****p < 0.0001, *Tonic* vs. *Onset*, p = 0.01, *Phasic* vs. *Random*, **p = 0.001). Spontaneous firing is given as an overall percentage, with exact number of neurons firing at rest over the total number of that type of cell shown in parentheses.

|  | **Phasic** | **Tonic** | **Onset** | **Random** |
| --- | --- | --- | --- | --- |
| **Resting Membrane Potential (mV)** | $-55 \pm 0.6$ | $-62 \pm 0.9$ | $-52 \pm 1.2$ | $-57 \pm 1.7$ |
| **Membrane Resistance ($G\Omega$)** | $2.4 \pm 0.3$ | $1.6 \pm 0.2$ | $1.8 \pm 0.4$ | $0.8 \pm 0.1$ |
| **Capacitance (pF)** | $42 \pm 1.4$ | $67 \pm 3.5$ | $49 \pm 4.6$ | $64 \pm 6.8$ |
| **Spontaneous Firing** | 69%<br>(71/103) | 92%<br>(71/77) | 0%<br>(0/23) | 37%<br>(7/20) |

### Intrinsic firing patterns and membrane properties of *Phasic* and *Tonic* neurons

Comparing the intrinsic membrane properties and firing patterns of the two most prevalent neuron types represented in our dataset underscored substantial differences in spontaneous activity, firing pattern, and firing frequency in response to stimulation. The majority of *Phasic* neurons were spontaneously active at the resting membrane potential (71/103, 69%; Table 1), and either fired tonic, evenly spaced action potentials, or shorter, bursts of action potentials with pauses of variable duration (Fig. 2A). *Phasic* neurons increased in firing frequency from an average of 4 Hz, in the spontaneously active neurons, up to 41 Hz in response to the maximal current injection (Fig. 2B); however, the firing duration decreased and action potential amplitudes became attenuated with increasing depolarizing steps (Fig. 2C). The decrease in firing duration and attenuation in action potential amplitude were also observed with repeated shorter (200 ms) current steps of 60 pA, 120 pA, and 180 pA (data not shown). The other major type of neuron based on intrinsic firing pattern was *Tonic* neurons which were spontaneously active at an average of 4 Hz (71/77, 92%; Table 1) and reached a maximal firing rate of 20 Hz in response to the 120 pA step (Fig. 2A,B). In addition to firing frequency, *Tonic* neurons were defined by stable firing for the full duration of each current step (Fig. 2C), which was also observed in response to shorter, repeated current steps (data not shown). *Tonic* neurons exhibited significantly slower firing frequency in response to depolarization compared to *Phasic* neurons (Two-way ANOVA, Main effect of neuron type, F_(1,178)_ = 70, p < 0.0001).

**Figure 2.**
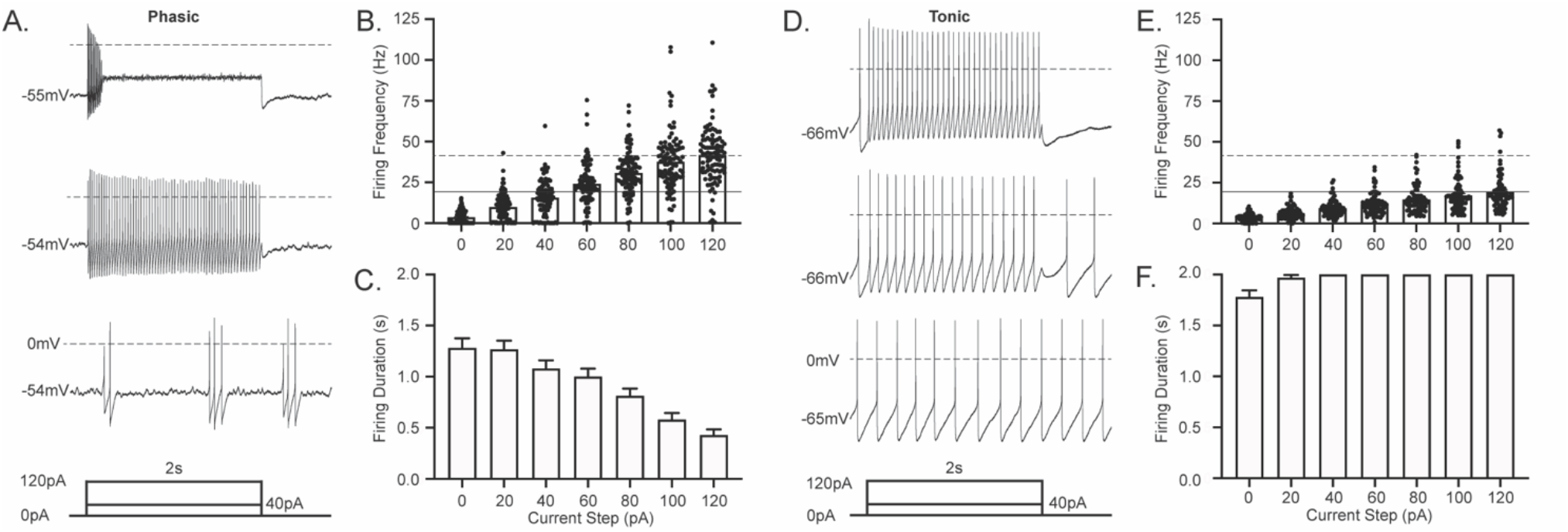
Defining features of the two most common vlPAG neurons. **A**. Representative trace of a recording from a *Phasic* vlPAG neuron from a naïve rat in response to 0 pA (bottom trace), 40 pA (middle trace), and 120 pA (top trace) current injections. Current injections were 2 s long, followed by 250 ms return to baseline (current protocol schematic below the traces). **B**. Firing frequency of all *Phasic* neurons for current steps ranging from 0 pA to 120 pA in 20 pA increments (n = 103). Dashed line shows maximal firing frequency at 120 pA for *Phasic* neurons, whereas solid line shows maximal firing frequency at 120 pA for *Tonic* neurons. **C**. Total firing duration of all *Phasic* neurons throughout each of the 2 s depolarizing current steps (n = 103). **D**. Representative trace of a recording from a *Tonic* vlPAG neuron from a naïve rat in response to 0 pA (bottom trace), 40 pA (middle trace), and 120 pA (top trace) current injections. Current injections were 2 s long after a 50 ms delay, followed by 250 ms return to baseline (as shown in the current protocol schematic below the traces). **E**. Firing frequency of all *Tonic* neurons for current steps ranging from 0 pA to 120 pA in 20 pA increments (n = 77). **F**. Compiled data showing total firing duration of all *Tonic* neurons throughout each of the 2 s depolarizing current steps (n = 77).

To determine whether the firing pattern and frequency were intrinsic, and independent of inhibitory and excitatory synaptic inputs, we used AMPA- and GABA_A_-receptor blockers (NBQX, 5 μM and gabazine, 10 μM) to block the synaptic inputs onto vlPAG neurons. Both *Phasic* and *Tonic* neurons maintain the same firing patterns and firing frequency at rest and in response to a series of depolarizing current steps in the absence of synaptic inputs (Fig. 3). Collectively, this suggests that the vlPAG has 2 main neuron types based on intrinsic membrane properties that exist in comparable proportions.

**Figure 3.**
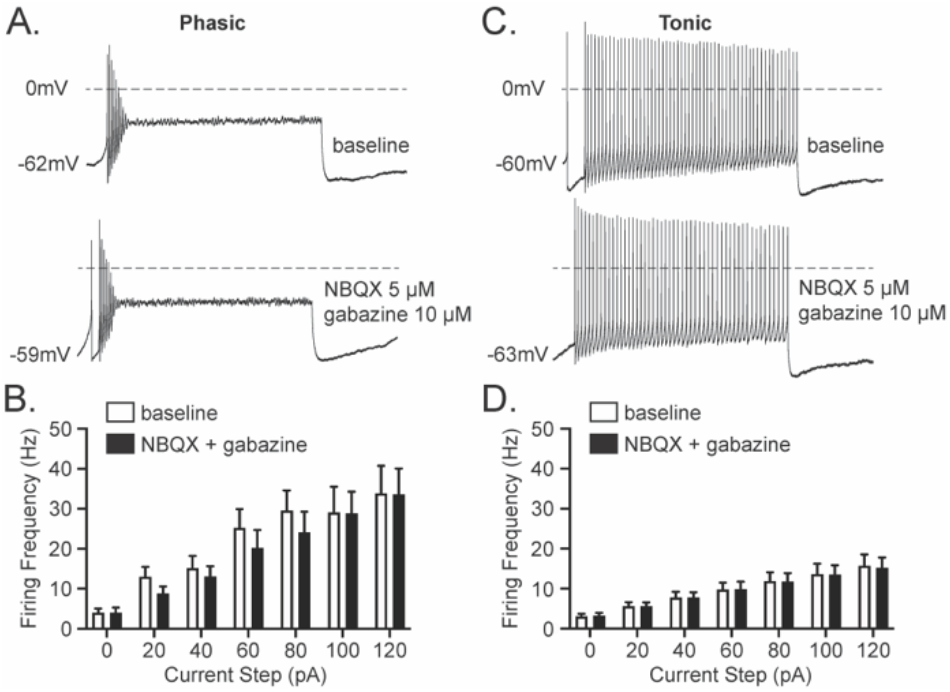
The firing patterns of *Phasic* and *Tonic* neurons are maintained in the absence of synaptic inputs. **A**. Representative traces of a recording from a *Phasic* vlPAG neuron from a naïve rat in response to 120 pA current injections before (top trace) and after synaptic blockers NBQX (5 μM) and gabazine (10 μM) (bottom trace). **B**. Compiled data showing firing frequency of all *Phasic* neurons for current steps ranging from 0 pA to 120 pA in 20 pA increments before and after synaptic blockers (Two-way Repeated Measures ANOVA, Effect of synaptic blockers, F_(1, 9)_ = 1.3, p = 0.3; n = 10). **C**. Representative traces of a recording from a *Tonic* vlPAG neuron from a naïve rat in response to 120 pA current injections before (bottom trace) and after synaptic blockers (top trace). **D**. Compiled data showing firing frequency of all *Tonic* neurons for all current steps before and after synaptic blockers (Two-way Repeated Measures ANOVA, Effect of synaptic blockers, F_(1, 10)_ = 0.002, p = 1.0; n = 11).

### MOR-mediated GIRK currents are observed within all 4 vlPAG neuron types

To determine whether postsynaptic MORs are selectively expressed on a single neuron type in the vlPAG, the MOR agonist DAMGO (5 μM) was used to elicit GIRK currents after evaluating the firing patterns of each neuron (Fig. 4A,B). Opioid-mediated GIRK currents were elicited in some, but not all, of the *Phasic, Tonic*, and *Onset* (Fig. 4C). All 3 *Random* neurons were opioid-sensitive but the small sample may have precluded observing opioid-insensitive neurons in this group. Opioid-insensitive neurons exhibited GABA_B_-mediated GIRK currents, indicating that these neurons express GIRK channels and not MOR’s, as we noted in our previous study (McPherson et al., 2018). Average GIRK currents in the *Phasic, Tonic*, and *Onset* neurons were ∼10 pA (Fig. 4D). Loose-patch recordings demonstrated clear DAMGO-mediated inhibition of spontaneous firing in naïve rats (naïve = 66% ± 5%, n = 56) indicating that the small GIRK currents were sufficient to inhibit firing. Mixed postsynaptic MOR expression within each neuron type motivated the search for additional firing features that indicate opioid sensitivity of a subpopulation of neurons.

**Figure 4.**
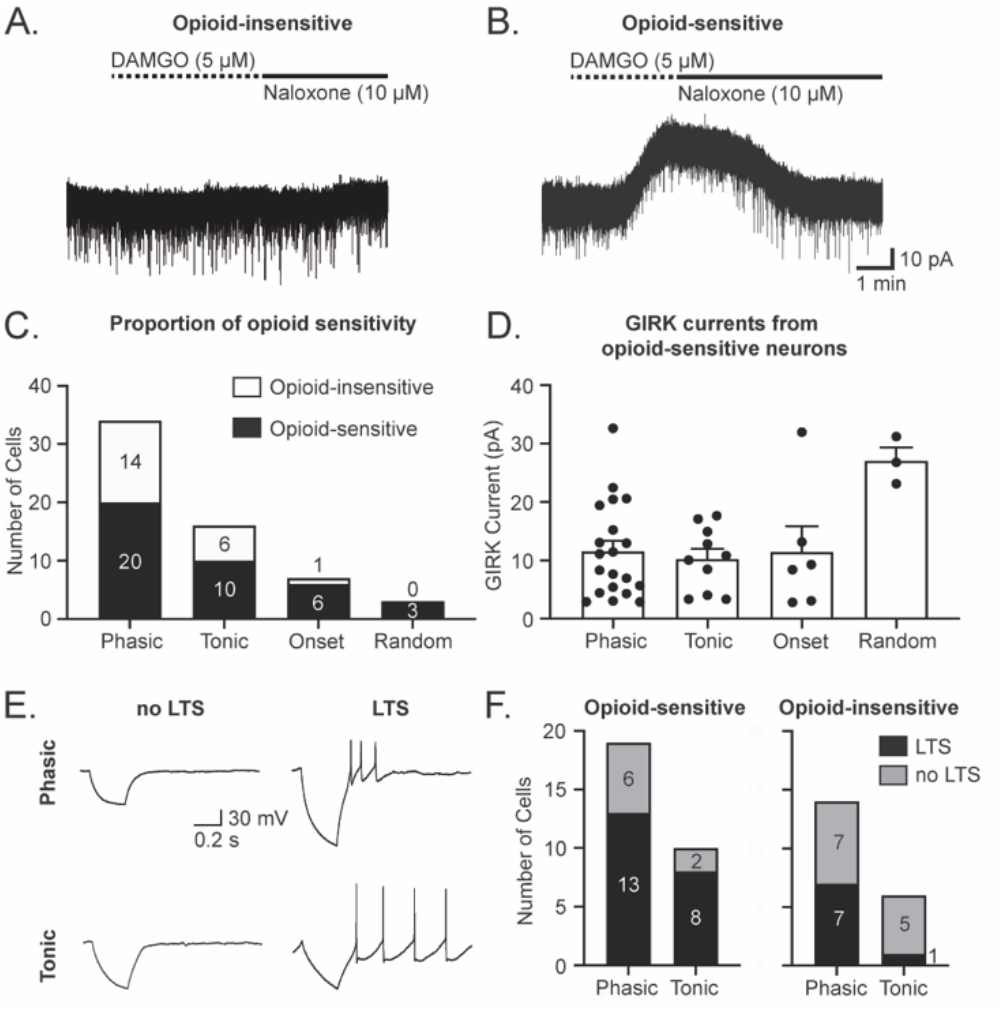
Opioid sensitivity is not restricted to selective neuron types in the vlPAG. **A**. Representative trace from a vlPAG neuron in which the selective MOR agonist DAMGO (5 μM) did not elicit an outward current (opioid-insensitive). **B**. Representative trace from a vlPAG neuron in which DAMGO (5 μM) elicited an outward current that was reversed by the MOR antagonist naloxone (10 μM) (opioid-sensitive). **C**. Number of neurons exhibiting DAMGO-mediated GIRK current responses (opioid-sensitive) compared to those that did not (opioid-insensitive) in each neuron type (n = 60). **D**. DAMGO-mediated GIRK current amplitude (pA) data for opioid-sensitive neurons in each neuron type (One-way ANOVA, main effect of neuron type, F_(3, 35)_ = 3.9, p = 0.02; Tukey’s post-hoc test *Phasic vs. Random* p = 0.01, *Tonic vs. Random* p = 0.01, and *Onset vs. Random* p = 0.04; n = 39). **E**. Sample whole-cell traces in response to 500 ms long -50 pA hyperpolarizing current steps, demonstrating neurons with and without low threshold spikes (LTS) in both *Phasic* and *Tonic* neuron populations. **F**. The Proportion of LTS and non-LTS neurons in *Phasic* and *Tonic* neuron groups, both in opioid-sensitive (left) and opioid-insensitive (right) groups.

Neurons exhibiting low-threshold spikes (LTS) have been previously described to be correlated with opioid sensitivity in mouse vlPAG. We found that both *Phasic* and *Tonic* groups had neurons with and without LTS in response to hyperpolarizing steps (Fig. 4E). More specifically, 6/19 opioid-sensitive *Phasic* neurons and 2/10 opioid-sensitive *Tonic neurons* showed no LTS, whereas in opioid-insensitive neurons, 7/14 *Phasic* and 1/6 *Tonic* neurons had LTS (Fig. 4F). Thus, LTS activity was not a robust indicator of opioid sensitivity in the rat vlPAG.

### Persistent inflammation selectively enhances spontaneous activity in *Phasic* neurons

Defining intrinsic firing properties in naïve rats provided the framework to evaluate alterations to intrinsic properties after acute and persistent inflammation. Both acute (2 h) and persistent inflammation (6 d) induce Fos expression in the vlPAG (Fig. 5A,B). To determine the underlying activity that could be driving this enhancement in Fos expression we examined the spontaneous firing frequency of *Phasic* and *Tonic* neurons after acute (2 h) and persistent (5-7 d) inflammation. *Tonic* neurons remained unchanged across both post-CFA time points (Fig. 5C), while the spontaneous firing frequencies of *Phasic* neurons doubled after persistent inflammation compared to *Phasic* vlPAG neurons in naïve rats and rats 2 h post-CFA (Fig. 5D). Importantly, the proportion of these 4 distinct neuron types were the same between the naïve condition (n = 223) and after acute (n = 64) and persistent (n = 84) inflammation (chi-square = 12.44, df = 6, p = 0.053), suggesting that there were no firing pattern changes induced by inflammation.

**Figure 5.**
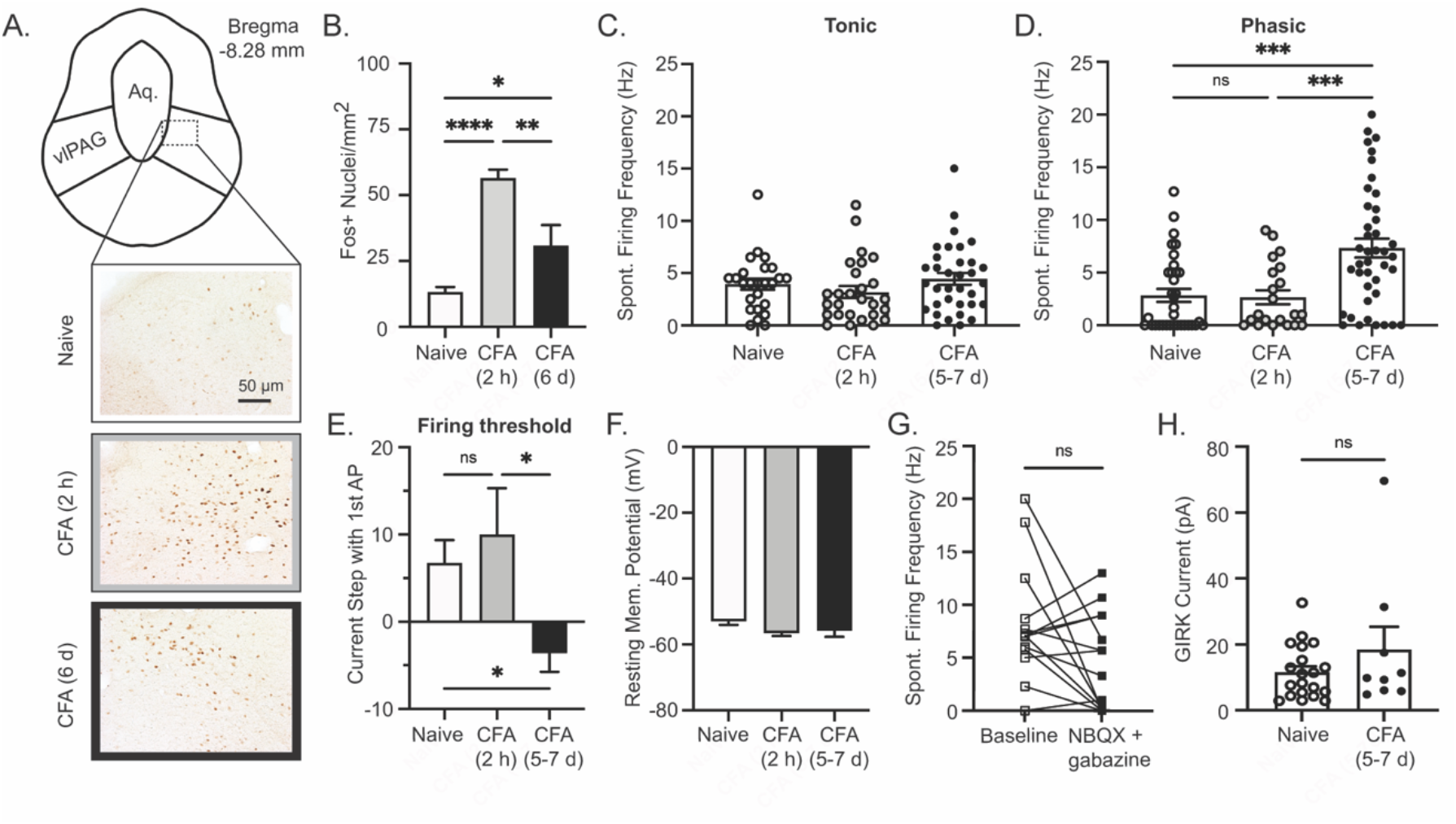
Persistent inflammation increases firing and decreases firing thresholds of *Phasic* neurons. **A**. Region of interest (vlPAG) outlined and representative image location demonstrated with dashed box. Representative images from Fos immunohistochemistry of tissue from naïve rats or rats sacrificed 2h or 6 d after CFA injection to the hind paw; cerebral aqueduct is labeled (Aq.), scale bar = 50 μm for all images. **B**. Average Fos+ nuclei/mm^2^ for naïve rats and those 2 h or 6 d after CFA injections (One-way ANOVA, F_(2, 11)_ = 26.3, p < 0.0001, Tukey’s post-hoc test, naïve vs CFA 2 h, ****p < 0.0001, naïve vs CFA 6 d, *p = 0.03, and CFA 2 h vs CFA 6 d, **p = 0.01). **C**. Spontaneous firing frequency of *Tonic* neurons after acute or persistent inflammation (One-way ANOVA, F_(2, 83)_ = 1.3, p = 0.3; naïve, n = 25, CFA, 2h n = 27, CFA 5-7 d, n = 34). **D**. Firing frequency of *Phasic* neurons shows significant increase in spontaneous firing frequency after persistent inflammation (One-way ANOVA, F_(2, 93)_ = 11.7, p < 0.0001; Tukey’s post-hoc test, naïve vs CFA 5-7 d, ***p = 0.0002 and CFA 2 h vs CFA 5-7 d, ***p = 0.001; naïve n = 34, CFA 2h n = 21, CFA 5-7 d n = 41) **E**. Firing thresholds (One-way ANOVA, F_(2, 91)_ = 5.6, p = 0.005, Tukey’s post-hoc test, naïve vs CFA 2 h, p = 0.8, naïve vs CFA 5-7 d, *p = 0.03, and CFA 2 h vs CFA 6 d, *p = 0.01). **F**. Resting membrane potential of *Phasic* neurons remained stable across all conditions (One-way ANOVA, F_(2, 90)_ = 1.4, p = 0.2). **G**. Spontaneous firing frequency of *Phasic* neurons before and after synaptic blockers in CFA (5-7 d) treated rats (Paired t-test, t _(14)_ = 1.9, p = 0.07). **H**. DAMGO-mediated GIRK current amplitude comparing naïve group (data from Fig. 4D) and after CFA (5-7 d) (Unpaired t-test, t_(27)_ = 1.3, p = 0.2).

To determine whether the increase in spontaneous firing of *Phasic* neurons was a result of changes to intrinsic firing properties, we looked at firing threshold, R_in_, RMP, and the firing response after removing synaptic inputs (using NBQX and gabazine as described previously). In naïve rats, 14/34 *Phasic* neurons required a depolarizing current step to fire, whereas after persistent inflammation most *Phasic* neurons fired spontaneously. Only 4/42 neurons required a depolarizing current step while 9/42 *Phasic* neurons fired during hyperpolarizing steps, resulting in a significant reduction in firing threshold after persistent inflammation (Fig. 5E). The RMP (Fig. 5F) and R_in_ (One way ANOVA, F_(2,60)_ = 0.1, p = 0.9) showed no changes across the two time points post-CFA injection. In addition, blockade of synaptic inputs did not affect the spontaneous firing frequencies of *Phasic* neurons (Fig. 5G). Collectively, these findings substantiate that the enhanced spontaneous firing rate of *Phasic* neurons is due to alterations to the intrinsic membrane properties.

DAMGO-mediated inhibition of spontaneous firing after persistent inflammation was comparable to that seen in naïve rats (naïve = 66% ± 5%, CFA (5-7 d) = 77% ± 5%; t_(91)_ = 1.5, p = 0.1, n = 56 naïve, n = 37 CFA (5-7 d)). Opioid-sensitive and -insensitive *Phasic* neurons were observed after persistent inflammation (20/34 opioid-sensitive neurons in naïve and 9/15 after CFA (5-7 d)), with comparable GIRK current amplitudes (Fig. 5H). There was no significant correlation between spontaneous firing frequency and DAMGO-mediated GIRK currents (simple linear regression, R^2^ = 0.05, p = 0.41). Thus, *Phasic* neurons in the vlPAG that are activated with persistent inflammation do not represent a homogenous MOR-expressing neuron type.

## Discussion

We examined intrinsic membrane properties of vlPAG neurons in naïve rats and in rats with acute (2 h) or persistent (5-7 d) inflammation. In all conditions we observed 4 types of neurons with distinct, intrinsic firing patterns. The firing patterns did not segregate with opioid sensitivity as MOR-activated GIRK currents were observed in a subpopulation of each neuron type. Despite robust Fos expression after acute inflammation, intrinsic firing properties remained unchanged until the persistent inflammation time point, where *Phasic* neurons displayed an increase in spontaneous firing frequency and lowered firing threshold. Further evaluation of the role of synaptic inputs after persistent inflammation confirmed that the enhanced activity of *Phasic* neurons was independent of synaptic inputs.

### Firing patterns and membrane properties define vlPAG neuron subtypes

We used voltage and current-clamp step protocols (Prescott and De Koninck, 2002; Sedlacek et al., 2007; Pradier et al., 2019) to initially characterize 223 vlPAG neurons from naïve animals. We observed 4 distinct firing patterns: *Phasic, Tonic, Onset*, and *Random. Phasic* and *Tonic* neurons comprise ∼80% of the neurons in the region and were evenly distributed across its rostral-caudal axis. Importantly, removal of synaptic inputs did not alter firing pattern or firing frequencies, demonstrating the intrinsic nature of the firing patterns. A previous study surveyed 33 neurons in the most caudal region of the mouse vlPAG, and observed the same firing patterns. Similarly, *Phasic, Tonic*, and *Onset* firing patterns are also represented in lamina I of the spinal dorsal horn, another key region in the processing and relaying of nociceptive information (Prescott and De Koninck, 2002). These similarities raise the question of whether neurons with these distinct firing patterns are functionally important for pain processing.

Given the key role the vlPAG plays in opioid-induced analgesia and the hypothesized postsynaptic MOR expression on GABAergic interneurons, we considered the possibility that opioid sensitivity may map onto these functionally distinct neuron types. MORs are expressed on a subpopulation of postsynaptic cell bodies within the vlPAG and are coupled to GIRKs, producing hyperpolarization (Chieng and Christie, 1994; McPherson et al., 2018). At least some of the MOR expressing neurons are GABAergic neurons with local connections (Barbaresi and Manfrini, 1988; Reichling and Basbaum, 1990; Chieng and Christie, 1994; Kalyuzhny and Wessendorf, 1998); however, opioids also inhibit a small proportion of RVM-projecting vlPAG neurons (Osborne et al., 1996). Consistent with these studies, approximately 22% of MOR-expressing neurons in the vlPAG are GABAergic, whereas 77% are glutamatergic (Zhang et al., 2020). These results indicate that neurotransmitter content (GABA vs. glutamate) alone is not a useful marker for vlPAG circuits. After determining the firing pattern of a neuron, we recorded GIRK current responses to DAMGO application, finding both opioid-sensitive and opioid-insensitive neurons within *Phasic, Tonic*, and *Onset* groups. A previous study observed opioid-mediated GIRK currents specifically in neurons exhibiting low-threshold spikes (LTS). DAMGO-mediated GIRK currents were observed in 5 neurons with LTS, with the remaining 4 neurons without LTS being opioid-insensitive (Park et al., 2010). Within our larger dataset in rats, LTS were observed in 50 *Phasic* and *Tonic* neurons and did not correlate with opioid sensitivity. These results further indicate that MOR expression is not a reliable marker of vlPAG neurons based on their intrinsic membrane properties.

Evaluation of the impact of opioids on spontaneous firing of vlPAG neurons using loose-patch recordings showed that MOR activation inhibited spontaneous firing in the majority of our experiments. Although we anticipated disinhibition of firing in some neurons (Lau et al., 2020; Morgan et al., 2020), disinhibition appears to be more prominent in RVM-projecting neurons that are not spontaneously active (Lau et al., 2020). Our loose-patch methods can only detect spontaneously active neurons, which make up 69% of *Phasic* and 92% of *Tonic* neurons, suggesting that opioid disinhibited neurons exist within the smaller population of quiescent neurons.

### Persistent inflammation increases spontaneous firing and decreases firing thresholds of *Phasic* neurons

We used CFA injections into the hindpaws of rats to produce persistent inflammation and were interested in whether intrinsic membrane properties were changed at either an acute (2 hr) or persistent time point (5-7 d). Fos, an immediate early gene upregulated in strongly activated neurons, displays peak expression 90-120 mins after a given stimuli. Fos is enhanced in the vlPAG in response to noxious stimuli (Keay and Bandler, 1993; Keay et al., 2000), and 25% of the Fos expressing vlPAG neurons following intramuscular formalin injections are RVM-projecting (Keay et al., 2000). In these studies, we did not introduce acute noxious stimulation prior to perfusing the brains so Fos staining represented ongoing activity of vlPAG neurons. Fos labeling indicated strong neuronal activation after acute (2 h) inflammation that was reduced by the persistent (5-7 d) time point, although still increased relative to naïve levels. To determine whether acute or persistent inflammation activates vlPAG neurons broadly or activates a subset of neurons, we assessed the same intrinsic firing properties protocols used in naïve rats. Importantly, in the CFA-treated rats, vlPAG neurons displayed all 4 firing patterns in similar proportions, suggesting no inflammation-induced changes in firing patterns.

In contrast to the Fos data, persistent inflammation increased spontaneous firing of *Phasic* neurons, which was not observed at the acute inflammation time point—suggesting that this was a time-dependent adaptation. Thus, the more robust Fos expression seen 2 h post-CFA injection is likely a result of an increase in nociceptive inputs with the development of inflammation. In the RVM, a target region for vlPAG afferents, CFA induces increased firing for the first 30 mins before returning to baseline (Cleary and Heinricher, 2013). Increased Fos activation in CFA-treated rats was maintained in the vlPAG until 5-7 d post-CFA and increased spontaneous firing of Phasic neurons became evident at this time point. *Tonic* neurons exhibited no changes in firing activity after acute or persistent inflammation, reinforcing the selective activation of the *Phasic* neurons. The activated *Phasic* neurons in the vlPAG may represent an ensemble of neurons (Whitaker et al., 2016; Whitaker et al., 2017; Whitaker and Hope, 2018) encoding continued peripheral nociceptor activation or inflammatory processing through alterations in intrinsic membrane properties, such as enhanced spontaneous activity and lowered firing threshold.

### Relevance to descending modulation of pain

Neurons within the vlPAG are known to have differential firing responses to acute noxious stimuli *in vivo* (Heinricher et al., 1987; Tryon et al., 2016), however, alterations to firing in persistent pain states and the resulting consequence on the descending pain modulatory circuit have not been fully defined. The vlPAG has been studied primarily in the context of descending *inhibition* of pain (Basbaum and Fields, 1984; Morgan et al., 1991; Vanegas and Schaible, 2004). However, a growing number of studies are providing evidence that the vlPAG plays a role in facilitation of pain (Heinricher et al., 2004; Dubový et al., 2018; Ni et al., 2019). In addition, neuroimaging in humans has implicated the PAG in central sensitization (Zambreanu et al., 2005). In the case of neuropathic pain induced by paclitaxel, innocuous stimuli increase firing of a subpopulation of vlPAG neurons in *in vivo* recordings (Samineni et al., 2017). These neurons fire with burst firing suggesting that they may represent the *Phasic* neurons in this study. Chronic constriction injury (CCI) increases firing frequency of some vlPAG neurons *in vitro* as well (Du et al., 2013).

An adaptation observed with both inflammation and neuropathic pain is an increase in spontaneous GABA release in the vlPAG (Hahm et al., 2011; Tonsfeldt et al., 2016). Although it is not clear whether this enhanced GABA tone originates from vlPAG afferents coming from other brain regions or vlPAG GABA neurons that send local collaterals, local TTX infusion within the vlPAG decreased extracellular GABA levels detected with microdialysis, indicating local GABAergic neurons contribute to vlPAG GABA tone (Maione et al., 1999). Increased GABA tone would contribute to descending pain facilitation and reduced morphine-mediated analgesia, similar to the effects of muscimol microinfusions into the vlPAG (Zambotti et al., 1982), by altering engagement with and subsequent activity of downstream RVM neurons (Moreau and Fields, 1986). It is possible that *Phasic* neurons are GABA neurons with local collaterals, so that enhanced spontaneous firing after persistent inflammation contributes to GABA tone.

In summary, this study characterized the intrinsic membrane properties of a large number of vlPAG neurons, identifying 4 distinct neuron types with mixed opioid sensitivity. Using this framework for defining neuron types we determined that persistent inflammation increases the spontaneous activity and lowers the firing threshold of *Phasic* neurons, in the absence of synaptic inputs. Altogether, this study increases our understanding of neuronal diversity within the vlPAG and promotes further investigation on the potential role of enhanced spontaneous firing of *Phasic* neurons in the bidirectional control of inflammatory pain.

## Acknowledgements

The studies were supported by the National Institute of Drug Abuse (NIDA) 1 R01 DA042565 (SLI) and 1 F31 DA052114-01 (CAB). KBM was supported by the National Science Foundation Graduate Research Fellowship Program. We thank Dr. Mary Heinricher for critical reading of the manuscript.

